# Architecture and topologies of gene regulatory networks associated with breast cancer, adjacent normal, and normal tissues

**DOI:** 10.1101/2022.10.10.511680

**Authors:** Swapnil Kumar, Vaibhav Vindal

## Abstract

Most cancer studies employ adjacent normal tissues to tumors (ANTs) as controls, which are not completely normal and represent a pre-cancerous state. However, the regulatory landscape of ANTs and how it differs from tumor and non-tumor-bearing normal tissues is largely unexplored. Among cancers, breast cancer is the most commonly diagnosed cancer and a leading cause of death in women worldwide, with a lack of sufficient treatment regimens due to various reasons. Hence, we aimed to gain deeper insights into normal, pre-cancerous, and cancerous regulatory systems of the breast tissues towards the identification of ANT and subtype-specific candidate genes. For this, we constructed and analyzed eight gene regulatory networks (GRNs), including five different subtypes (*viz.* Basal, Her2, LuminalA, LuminalB, and Normal-Like), one ANT, and two normal tissue networks. Whereas several topological properties of these GRNs enabled us to identify tumor-related features of ANT; escape velocity centrality (EVC+) identified 24 functionally significant common genes, including well-known genes such as *E2F1, FOXA1, JUN, BRCA1, GATA3, ERBB2,* and *ERBB3* across different subtypes and ANT. Similarly, the EVC+ also helped us to identify tissue-specific key genes (Basal: 18, Her2: 6, LuminalA: 5, LuminalB: 5, Normal-Like: 2, and ANT: 7). Additionally, differential correlation along with functional, pathway, and disease annotations highlighted the cancer-associated role of these genes. In a nutshell, the present study revealed ANT and subtype-specific regulatory features and key candidate genes which can be explored further using *in vitro* and *in vivo* experiments for better and effective disease management at an early stage.

## Introduction

Adjacent normal tissues to tumors (ANTs) are widely used as controls in various cancer studies. However, a comprehensive study on ANTs by Slaughter et al. has revealed the presence of an acquired step-by-step cumulative carcinogenic process comprising various genetic alterations, thus portraying the ANTs as a pre-neoplastic, intermediate state composed of morphologically normal but altered cells at the molecular level (Slaughter et al. 1953). Another study on ANTs has also demonstrated the differentiating characteristics of ANTs from healthy tissues (Aran et al. 2017). In spite of this, the regulatory landscape of ANTs is not much explored and hence lacks substantial knowledge about how it differs from the regulatory landscapes of tumor and non-tumor-bearing normal tissues. In-depth knowledge of the comparative regulatory landscape of ANTs with different subtypes and normal tissues can help to understand the early or pre-cancerous events in a better way which leads to the development and progression of the disease. This may further lead to identifying more specific and novel candidate genes at the subtype level to improve the treatment decisions for an early and more effective disease management.

Breast cancer is the leading cause of death (15.5%) and also the most commonly diagnosed cancer (24.5%) in women worldwide, as per the estimated data of GLOBOCAN 2020 (Sung et al. 2021). Even with the availability of various treatments such as hormone therapy, chemotherapy, radiotherapy, targeted therapy, and surgery, it is still the leading cause of concern. For example, patients diagnosed with early-stage tumors in about 30% of cases, even after removing the tumors, are prone to recurrence in distant organs (McAllister et al. 2008). Additionally, those diagnosed with late-stage and distant metastases are incurable (Redig and McAllister, 2013).

Breast cancer is a heterogeneous disease with multiple subtypes. It can be stratified into five different molecular subtypes based on the expression profiles of hormone receptors such as estrogen/progesterone receptor (ER/PR) and human epidermal growth factor receptor 2 (HER2). The five subtypes include Basal-like (ER-/PR-and HER2-), HER2-positive (ER+/PR+ and HER2+), Luminal A (ER+/PR+, HER2- and Ki67 low), Luminal B (ER+/PR+, HER2- or HER2+, and Ki67 high), and Normal-like (ER+/PR+, HER2-, and Ki67 low) (Dai et al. 2015). These molecular subtypes of breast cancer exhibit varying biological features and hence respond differently to treatment processes leading to disparate clinical outcomes. Further, these also occur at different rates and are more or less aggressive in nature, with variations in long-term survival rates (Yersal and Barutca, 2014). Despite the substantial advancement in the treatment process of breast cancer, to date, a proper treatment at the level of its different subtypes is still limited in terms of the lack of precise molecular targets for each one of them (Meng et al. 2016). In this regard, some progress has been made toward identifying subtype-specific as well as metastatic tissue-specific molecular targets or biomarkers (Calza et al. 2006; Walker et al. 2007; Shan et al. 2014; de Azevedo et al. 2023). However, these are still inadequate to efficiently tackle subtype-specificity-based treatments’ limitations.

The network theory-based approach allows us to investigate the vast amount of high-throughput data in a novel way (Srivastava et al. 2014; Shi et al. 2017; Ruiz Amores and Martínez-Antonio 2022; Kumar et al. 2023a; Zhou et al. 2023). Genes and their products work together by interacting with each other to perform various essential cellular functions and processes. These genes, by interacting with each other, form a complex network. A diseased cell, including a cancerous cell, can be represented as a gene regulatory network (GRN) or a protein-protein interactions network (PPIN). The GRNs are very important for understanding the regulatory mechanism of genes and, thus, gene expression heterogeneity across normal and diseased cells. A comprehensive analysis of GRN representing cancerous cells can provide valuable insight into its behavior and may lead to the discovery of new and efficient biomarkers. Hence, the detailed exploration of the GRN of ANTs and its comparison with the GRNs of different breast cancer subtypes can help to identify key candidate genes that can be considered as early-stage-specific genes. These early-stage-specific candidate genes can be explored and utilized further to develop more efficient, novel biomarkers and therapeutics for the disease.

In the present study, first, we identified differentially expressed genes (DEGs) across different subtypes of breast cancer and adjacent normal tissues using gene expression profiles of the tumor, adjacent normal, and normal tissues of the human breast. It followed the construction and subsequent analysis of GRNs of each subtype, ANT, and normal breast tissue (NBT) using the regulatory information of transcription factors (TFs) and target genes available in various public databases. Furthermore, the TF-gene regulatory information for these tissue types was also predicted using expression profiles of DEGs along with prior data such as PPI and motif. Besides, all DEGs of each subtype were also analyzed using differential correlation, disease annotation, and functional enrichment analysis. The differential correlation approach has been used previously to identify differentiating features between two conditions or subtypes of a disease (Fukushima 2013; Zhou et al. 2021; Kumar et al. 2023a). This comprehensive study helped us identify breast cancer’s condition and subtype-specific features. The structural properties of GRNs also enabled us to identify some functionally significant genes responsible for the development and progression of the disease. These important genes can be explored further using *in vitro* and *in vivo* experiments to develop a more accurate and effective treatment strategy for better disease management.

## Methods

An experimental overview of the current study is given in Figure 1.

**Fig 1.**
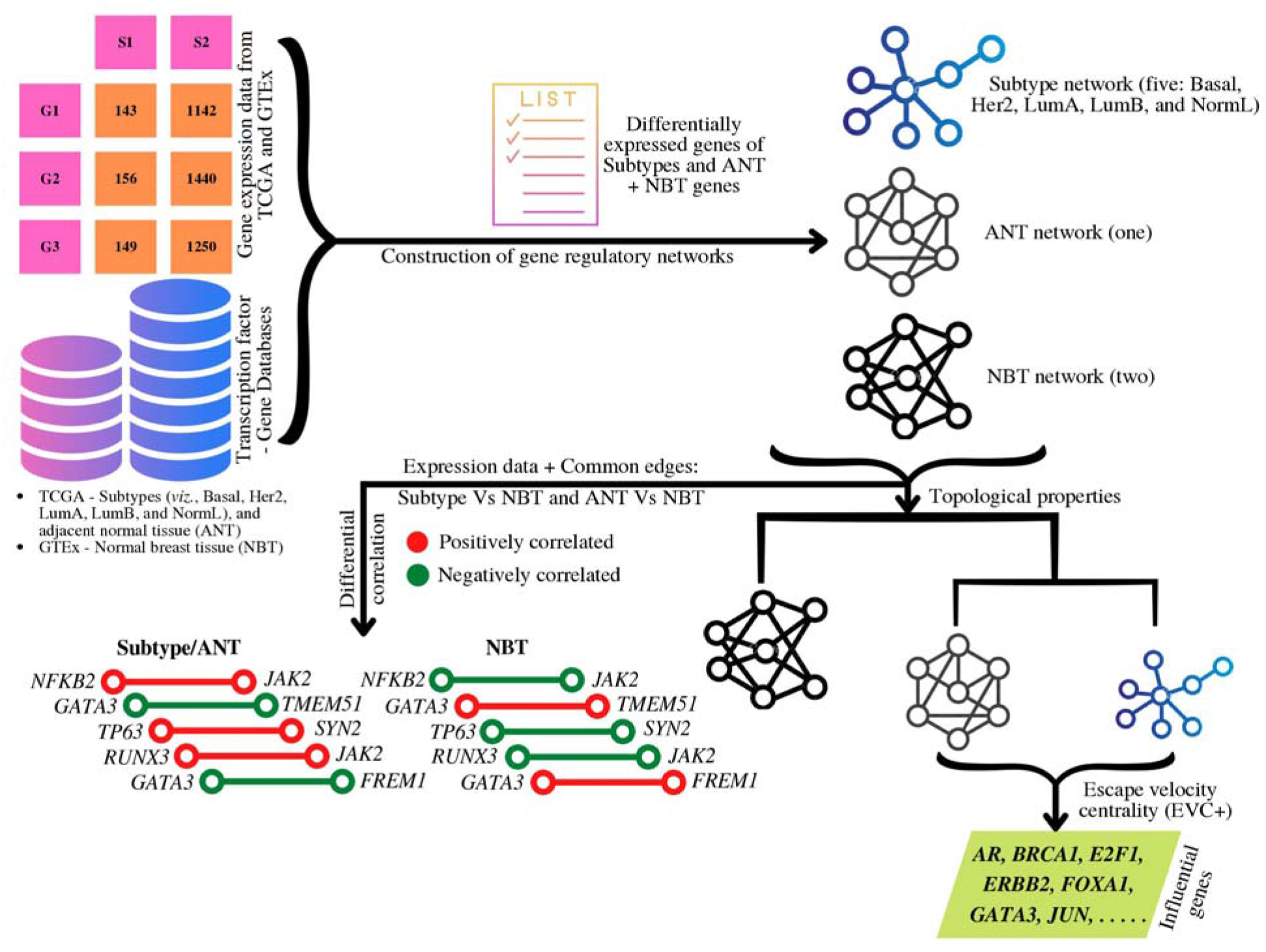
Experimental overview of the study.

### Retrieval of gene expression data

Gene expression profiles of breast tumor tissues and adjacent normal tissues (ANT) from The Cancer Genome Atlas (TCGA) project (Weinstein et al. 2013; Tomczak et al. 2015), along with gene expression profiles of normal breast tissues (NBT) from the Genotype-Tissue Expression (GTEx) database (Lonsdale et al. 2013), were accessed through the recount2 project (Collado-Torres et al. 2017). The recount2 project provides uniformly processed and quantified RNA-seq data derived from the GTEx, Sequence Read Archive (SRA), and TCGA projects for various downstream analyses, including differential expression analysis. All these three datasets were pre-processed and normalized for further use using the TCGAbiolinks R/Bioconductor package (Colaprico et al. 2016).

The gene expression data of tumor tissues were further subdivided or stratified into expression profiles for five different subtypes of breast cancer, *viz.,* Basal, Her2, Luminal A (LumA), Luminal B (LumB), and Normal-like (NormL), based on the sample’s information as available from the Genomics Data Commons (GDC) portal of TCGA.

### Acquisition of breast tissue genes

Breast tissue genes were also collected, first, by processing the gene expression data of the NBT and termed as gene set one (GS1). For this, genes having mean values of transcripts per million (TPM) and counts per million (CPM) equal to or more than one across all samples were considered as breast tissue genes. The TPM and CPM values of genes were calculated with the help of the “convertCounts” function available in the DGEobj.utils R package. Secondly, a set of breast tissue genes were also retrieved from The Human Protein Atlas (Uhlén et al. 2015) and were referred to as the gene set two (GS2). These two sets of breast tissue genes (GS1 and GS2) will further be used for the construction of two GRNs of the NBT.

### Identification of differentially expressed genes

To identify differentially expressed genes (DEGs) in each subtype, gene expression data of all five subtypes were compared with gene expression profiles of NBT using the DESeq2 (Love et al. 2014), edgeR (Robinson et al. 2010), and limma R package (Ritchie et al. 2015). Similarly, DEGs of ANT compared to the normal tissues (NBT), were also identified. Thus, it resulted in six lists of DEGs (significantly up and down-regulated genes) across six different conditions. Only significantly differentially expressed genes (up and down) as per any two tools out of three were considered for further analysis. The expression profiles of the DEGs of different subtypes and ANT were extracted from the corresponding tissue data for subsequent analysis. Further, the expression data of these DEGs from normal tissues were also extracted for the downstream analysis.

### Construction of subtype, adjacent normal, and normal networks

The DEGs of each of five subtypes (Basal, Her2, LumA, LumB, and NormL), along with DEGs of ANT, were used to construct six GRNs (five subtypes and one ANT) with the help of seven different databases *viz.,* GRAND (Ben et al. 2022), GRNdb (Fang et al. 2021), hTFtarget (Zhang et al. 2020), ICON (Clauset et al. 2016), ORTI (Vafaee et al. 2016), RegNetwork (Liu et al. 2015), and TRRUSTv2 (Han et al. 2018). These databases store predicted as well as experimentally determined information on transcription factor (TF)-target genes. The GRAND database hosts several predicted GRNs derived from TF-gene information of various cell lines, tumor tissues, normal tissues, and others. Out of these GRAND GRNs, only TF-gene regulations predicted using the gene expression data of breast tumor tissues from TCGA and of NBT from GTEx were considered in the current study. These GRAND regulatory data had more than one million TF-gene regulations; therefore, only the top 10% of TF-gene regulations based on the edge score > 0 were used to identify regulations of DEGs for subtypes and of breast tissue genes for NBT network constructions from the TF-gene regulations of the breast tumor tissues (TCGA) and NBT (GTEx), respectively. The GS1 and GS2 lists of breast tissue genes as retrieved previously were also mapped to these seven databases to construct two NBT networks. Further, there was no TF-gene regulatory information on ANT in the GRAND database; therefore, the gene expression data of 19,061 protein-coding genes across 111 samples from ANT was used to predict TF-gene regulations with the help of the PANDA R package (Schlauch et al. 2017). From the predicted TF-gene regulations of ANT, only the top 10% of TF-gene regulations based on the edge score > 0 were considered to identify regulations of DEGs of ANT for the corresponding network construction.

### Regulatory network analysis

All five subtype networks (Basal, Her2, LumA, LumB, and NormL) and one ANT network along with two normal breast networks (NBT1 and NBT2) were analyzed using the igraph R package (Csardi and Nepusz, 2006) by assessing various topological properties. The topological properties considered for the present analysis were degree, clustering coefficient, betweenness centrality, average path length, average local efficiency, diameter, eigen centrality, and heterogeneity. Further, eight random networks (one for each original network) were constructed with the same degree sequences present in the original networks. These random networks were used to analyze the structure and organization of corresponding real networks by comparing their different topological properties.

Degree: The total number of nodes directly connected with a node of interest is called the Degree of that node. It is the most basic property of a node participating in any network, and the fraction of nodes having degree value k is called Degree distribution, i.e., *P*(k). Highly connected genes (hubs) in biological networks are thought to play key roles in the behavioral organization of the networks (Albert et al. 2000; Jeong et al. 2001; Han et al. 2004; Carlson et al. 2006).

Betweenness centrality: The fraction of shortest paths between all the nodes’ pairs passing through a node of interest is defined as the Betweenness centrality of that node (Freeman 1977). Clustering coefficient: The ratio of the number of triangles a node of interest makes with its neighboring nodes and the total number of possible triangles that node can have is defined as the Clustering coefficient (Watts and Strogatz 1998; Ravasz et al. 2002).

Average path length: It is the average number of edges along the shortest paths for all possible node pairs in a network. It is also known as the Characteristic path length or the Average shortest path length (Watts and Strogatz 1998).

Average local efficiency: It is the arithmetic mean of local efficiencies of all the nodes present in a network; where the local efficiency of a node is the average of the reciprocal distances between all its neighbors after removing that node and considering all the remaining nodes of the network (Latora and Marchiori 2001; Vragović et al. 2005).

Diameter: It is the length of the longest “shortest path” connecting any two nodes present in the network. The diameter of a network describes its interconnectedness and the ability of nodes to communicate with other nodes in the network (Albert et al. 2000; Wasserman and Faust 1994).

Eigen centrality: It is the value of the eigenvector corresponding to the largest eigenvalue of the adjacency matrix of a network, also known as eigenvector centrality (Bonacich 1972; Ruhnau 2000). It measures the influence of nodes in a network. If a node is connected to the high-scoring nodes, it has more contributions to the score of the node of interest compared to the equal number of connections with low-scoring nodes. A high eigenvector score of any node means that the node is connected with other nodes having high scores.

Heterogeneity: It is defined as the coefficient of variation of the connectivity distribution (Dong and Horvath 2007). All biological networks are very heterogeneous in nature; because some nodes are highly connected while the majority of nodes are less or very less connected.

### Network similarity assessment

Similarity among different GRNs in terms of nodes (representing genes) and edges (representing transcriptional regulations or interactions) points toward similarity in transcriptional activity or programs and thus similarity at the level of phenotypes. Thus, all five subtypes, one ANT, and two NBT networks were assessed for their similarity and dissimilarity with each other using *Jaccard* index (*J*-index). The *J*-index of networks can be calculated with the help of nodes and edges present therein. It is the ratio of the total number of common nodes or edges between two networks (intersection) to the total number of nodes or edges of both the networks (union). The value of *J*-index ranges between 0 to 1. If the value of *J*-index is 1, it indicates that two networks are 100% similar or identical; whereas if the *J*-index is 0, two networks have no similarity, i.e., the two networks are dissimilar to each other.

### Identification of influential genes

All subtype and ANT networks were analyzed to identify influential genes with the help of extended escape velocity centrality (EVC+) implemented in the NetVA R package (Kumar et al. 2023b). As proposed by Ullah et al., it is an extended version of the escape velocity centrality (EVC) which considers the number of connections a node can have along with its position in the network to identify key players (nodes) of the network. Thus, it provides a better ranking than any other, i.e., baseline network properties, including the degree, betweenness, and clustering coefficient of nodes based on the connections and positions of nodes participating in a network under study (Ullah et al. 2022).

The EVC+ of node ‘i’ can be defined as follows:

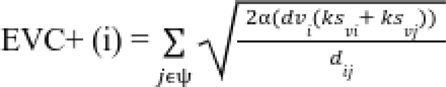

Where, α is a tunable factor which controls the degree effect, and its values range between 0.1 - 1, dv_i_ is the degree of node ‘i’, ks_vi_ and ks_vj_ are k-shell values of node ‘i’ and ‘j’ respectively, d_ij_ is the shortest path between two nodes ‘i’ and ‘j’ using Dijkstra algorithm, and jє_ψ_ is the set of neighbors of node ‘i’.

### Differential correlation analysis

The NBT1 network was compared with all five subtype networks and one ANT network one by one in a pairwise manner (i.e. Basal vs NBT1, Her2 vs NBT1, LumA vs NBT1, LumB vs NBT1, NormL vs NBT1, and ANT vs NBT1), and common edges between them were identified. Next, the expression profiles of DEGs of each subtypes (i.e. Basal, Her2, LumA, LumB, and NormL) from the corresponding tumor conditions (subtype tissues) along with the expression profiles of DEGs of ANT from the ANT itself were analyzed by comparing with the expression profiles of respective list of DEGs from the normal conditions (NBT) using the DiffCorr R package (Fukushima 2013). This resulted in a list of gene pairs (edges) significantly differentially correlated in case of different subtypes and ANT compared to the NBT. From the list of differentially correlated gene pairs of each subtype and also ANT, gene pairs showing switching behavior between the two conditions (tumor and NBT or ANT and NBT) were filtered out. Gene pairs with positive correlations in the tumor tissues or ANT, while negative correlations in the NBT and vice-versa were considered as gene pairs (edges) with switching behavior. Further, these switching gene pairs were filtered out based on their presence in the list of previously identified common edges between each subtype network and NBT1 network or ANT and NBT1 networks. These switching gene pairs or edges of all five subtypes and also ANT were further used to construct their corresponding networks and were visualized using the Cytoscape.

### Function, pathway, and disease enrichment analysis

All DEGs in the case of subtypes and ANT were further subjected to functional and pathway enrichment or over-representation analysis to decipher significantly over-represented terms *viz.* biological processes (BPs), molecular functions (MFs), and pathways (KEGG) associated with these genes. The functional and pathway enrichment analysis of DEGs was carried out using the clusterProfiler v4.4.4 (Wu et al. 2021) and visualized with the enrichplot R package (Yu 2022). The DEGs of subtypes and ANT were also checked for their associations with various diseases, including breast cancer. For this, the gene-disease associations using the disease-gene network (DGN) implemented in the DisGeNet database (Piñero et al. 2015) and the disease enrichments using the Disease Ontologies (DO) were inferred with the help of the clusterProfiler R package. The DisGeNet database hosts information on gene-disease associations as reported in previously published literature. All the enrichment terms with an adjusted *P*-value less than 0.05 were considered significant.

## Results

### Expression profiles of subtypes and normal tissues

The gene expression profiles of breast tumor tissues contained expression values for 56,002 genes across 1,210 samples. The expression profiles of normal breast tissues (NBT) contained expressions of 56,002 genes across only 92 samples. Among tumor tissue samples, there were 200 samples of Basal, 82 samples of Her2, 570 samples of LumA, 207 samples of LumB, 40 samples of NormL, and 111 samples of adjacent normal tissues (ANT). We considered only the expression profiles of 19090 protein-coding genes reported by the Human Gene Nomenclature Committee (HGNC) for the current study.

Before further analysis, these expression profiles were pre-processed and normalized, as discussed in the Methods section. After normalization, the data contained expression profiles of only 19079 genes in the case of tumors and normal tissues and 19060 genes in adjacent normal.

### Differentially expressed genes

The differentially expressed genes (DEGs) were identified with |FoldChange| > 1.5 and adj P-value < 0.05 across different subtypes, adjacent normal, and normal tissues using three commonly used R packages *viz.* DESeq2, edgeR, and limma. All subtypes and adjacent normal tissues were compared with normal breast tissues to identify DEGs. However, normal breast tissues were compared with all tumor tissues combined (including all subtypes) to identify DEGs across normal breast tissues. Thus, significantly up and down-regulated genes were selected for further analysis as per any two out of the three tools across all seven tissue types, including adjacent normal and normal tissues. The distributions of significantly up and down-regulated genes across all tumor and normal tissue samples are available in Table 1.

**Table 1.**
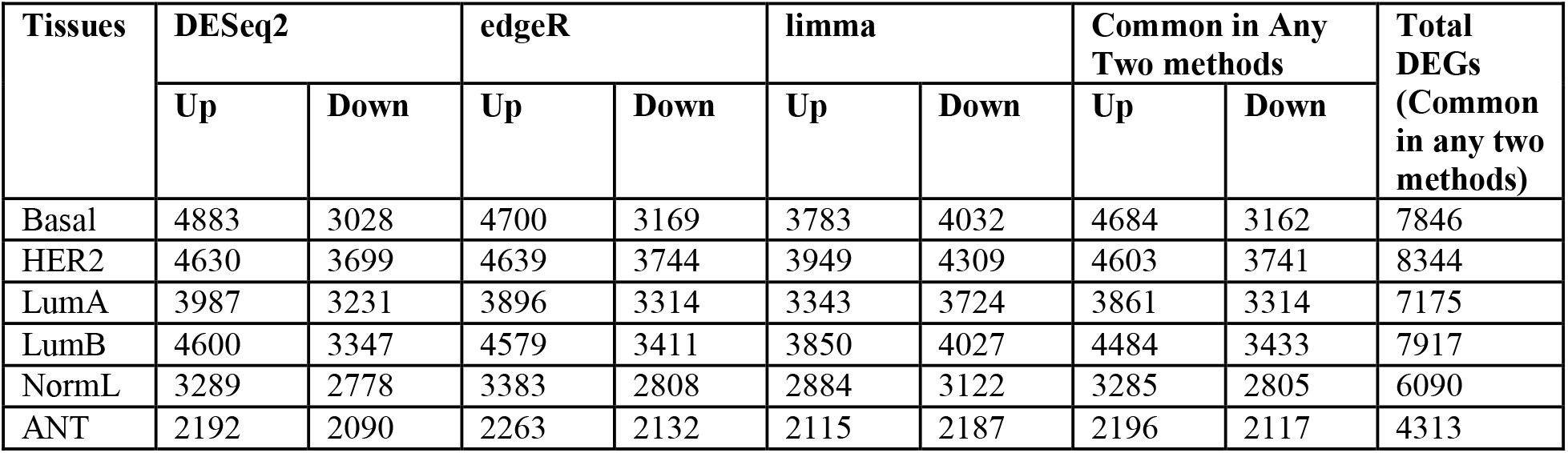
Distribution of significantly differentially expressed genes across all tissue samples with |FoldChange| > 1.5 and adj. P-value < 0.05.

Out of all DEGs in the Basal subtype, there were 792 genes unique to the Basal. Similarly, Her2, LumA, LumB, and NormL have 337, 150, 233, and 169 unique genes, respectively. Furthermore, the ANT also has 359 unique genes. These unique genes were present only in respective tissues but not in others. In addition to this, there were 1986 unique DEGs which were common among all five subtypes and ANT also. Similarly, a total of 3764 unique DEGs were common among all five subtypes.

### Gene regulatory networks of subtypes, adjacent normal, and normal tissues

Gene regulatory networks (GRN) for all five subtypes, adjacent normal and normal tissues were constructed by retrieving interactions data from five publicly available databases as discussed in the Methods section. The distributions of extracted edges across all five subtypes, ANT, and NBT are available in Table 2. The numbers of nodes and interactions present among them and in the largest connected components across all seven networks are described in Table 3.

**Table 2.** Distribution of edges collected from seven different sources for all seven tissue types.

| Network | GRAND | GRNdb | hTFtarget | ICON | ORTI | RegNetwork | TRRUST |
| --- | --- | --- | --- | --- | --- | --- | --- |
| Basal | 282347 | 47045 | 44031 | 1146 | 16713 | 34367 | 2419 |
| Her2 | 328704 | 46191 | 51656 | 1468 | 16960 | 40219 | 2735 |
| LumA | 254804 | 39753 | 41270 | 1117 | 18253 | 29956 | 2207 |
| LumB | 293235 | 44378 | 50848 | 1279 | 20067 | 38608 | 2610 |
| NormL | 168693 | 26279 | 29235 | 826 | 10827 | 18736 | 1577 |
| ANT | 30457 | 5547 | 18115 | 402 | 6589 | 9944 | 813 |
| NBT1 | 192533 | 82798 | 164055 | 3436 | 40483 | 119500 | 6247 |
| NBT2 | 183875 | 81421 | 157885 | 3376 | 43897 | 123603 | 6898 |

**Table 3.** Distribution of edges and vertices participating in the networks and largest connected components of the networks across all five subtypes, adjacent normal, and normal tissues.

| Network | N<sub>V</sub> | N<sub>E</sub> | N<sub>V</sub> (LCC) | N<sub>E</sub> (LCC) | alpha |
| --- | --- | --- | --- | --- | --- |
| Basal | 6100 | 32885 | 6099 | 32884 | 2.581 |
| Her2 | 6558 | 36240 | 6556 | 36238 | 2.564 |
| LumA | 5641 | 32941 | 5639 | 32939 | 2.708 |
| LumB | 6307 | 37652 | 6305 | 37650 | 2.628 |
| NormL | 4536 | 20455 | 4534 | 20453 | 2.598 |
| ANT | 2514 | 5826 | 2506 | 5820 | 2.887 |
| NBT1 | 11648 | 46519 | 11647 | 46518 | 1.759 |
| NBT2 | 11463 | 45952 | 11462 | 45951 | 1.769 |

Number of nodes, N<sub>V</sub>; Number of edges, N<sub>E</sub>; Largest connected component, LCC; Power law exponent of the degree distribution, alpha.

### Topological properties of networks

All networks, including ANT and NBT, were assessed based on different topological properties like degree, betweenness centrality, clustering coefficient, eigenvector centrality, average path length, average local efficiency, network diameter, and heterogeneity (Table 4). By analyzing these properties, we were able to draw some interesting inferences from these networks. All five subtypes and one ANT network follow the power law of degree distribution with alpha values between 2 to 3 (Table 3); however, both the NBT networks exhibit less than two, which may be due to the effect of the finite size observed in the case of other biological networks (Lotem et al. 2004). This proves that all the networks constructed in the present study are significant and represent true biological networks in terms of the structural organization of these networks, similar to other biological networks. The average degree values of all subtype networks are higher than the average degree values of normal network NBT. However, the average degree value of ANT is lower than that of all subtypes and NBT networks. Further, the average betweenness values of all five subtypes and one ANT network, *viz.,* Basal, Her2, LumA, LumB, NormL, and ANT, are lower than normal network NBT. However, the average betweenness value of the ANT network is the lowest among all networks. Next, all subtypes and ANT networks exhibited higher average values of clustering coefficient (<CC>) and eigenvector centrality (<EC>) than the normal network NBT. Moreover, the number of nodes with clustering coefficients equal to one and zero (CC = 1 and CC = 0) present in all five subtype networks was less than that of the normal network NBT. The number of nodes with CC = 1 and CC = 0 present in ANT was also less than that of the normal network NBT. Next, the values of average local efficiency (ALE) of all five subtype networks were higher than that of the normal networks (NBT1 and NBT2), whereas the ANT network had less value than both the normal networks. Thus, it indicated that the subtype networks were more efficient, and the ANT network was less efficient in cellular communication than the normal networks. Similarly, the average path lengths (APL) of all five subtype networks and one ANT network showed lower values than that of the normal networks (NBT1 and NBT2). Furthermore, the values of diameters of all five subtype networks and one ANT network were also less than that of the normal networks (NBT1 and NBT2). Next, the heterogeneity values of all five subtype networks and one ANT network were lower than that of the normal networks.

**Table 4.**
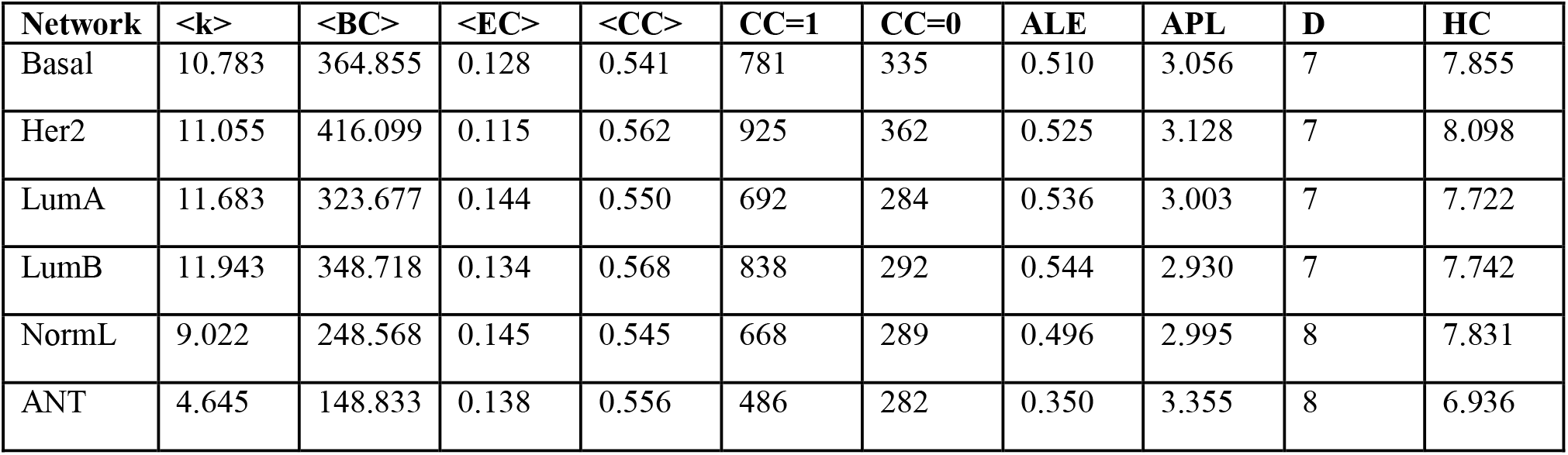
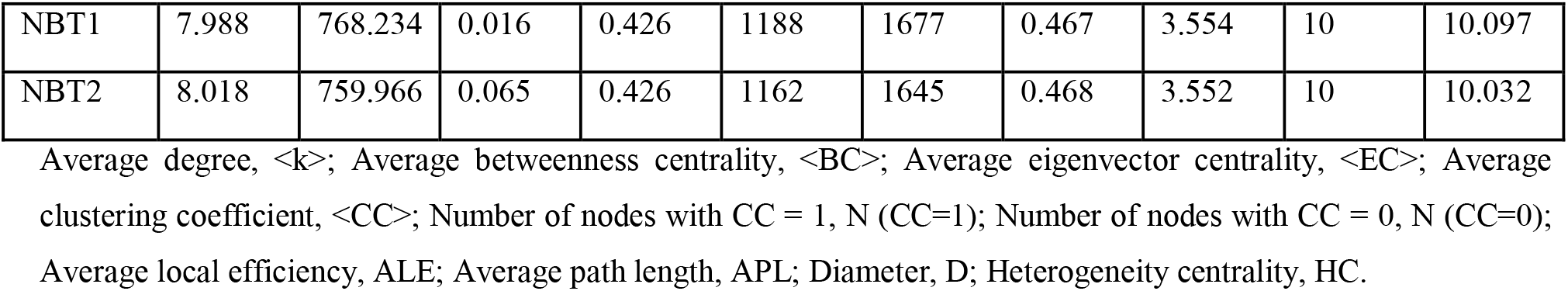
Various topological properties of all subtype and normal networks.

| Network | $\langle k \rangle$ | $\langle BC \rangle$ | $\langle EC \rangle$ | $\langle CC \rangle$ | CC=1 | CC=0 | ALE | APL | D | HC |
| --- | --- | --- | --- | --- | --- | --- | --- | --- | --- | --- |
| Basal | 10.783 | 364.855 | 0.128 | 0.541 | 781 | 335 | 0.510 | 3.056 | 7 | 7.855 |
| Her2 | 11.055 | 416.099 | 0.115 | 0.562 | 925 | 362 | 0.525 | 3.128 | 7 | 8.098 |
| LumA | 11.683 | 323.677 | 0.144 | 0.550 | 692 | 284 | 0.536 | 3.003 | 7 | 7.722 |
| LumB | 11.943 | 348.718 | 0.134 | 0.568 | 838 | 292 | 0.544 | 2.930 | 7 | 7.742 |
| NormL | 9.022 | 248.568 | 0.145 | 0.545 | 668 | 289 | 0.496 | 2.995 | 8 | 7.831 |
| ANT | 4.645 | 148.833 | 0.138 | 0.556 | 486 | 282 | 0.350 | 3.355 | 8 | 6.936 |
| NBT1 | 7.988 | 768.234 | 0.016 | 0.426 | 1188 | 1677 | 0.467 | 3.554 | 10 | 10.097 |
| NBT2 | 8.018 | 759.966 | 0.065 | 0.426 | 1162 | 1645 | 0.468 | 3.552 | 10 | 10.032 |

Average degree, $\langle k \rangle$ ; Average betweenness centrality, $\langle BC \rangle$ ; Average eigenvector centrality, $\langle EC \rangle$ ; Average clustering coefficient, $\langle CC \rangle$ ; Number of nodes with $CC = 1$ , $N(CC=1)$ ; Number of nodes with $CC = 0$ , $N(CC=0)$ ; Average local efficiency, ALE; Average path length, APL; Diameter, D; Heterogeneity centrality, HC.

When these properties of random networks (Table S1) were compared with those of the corresponding real networks, they were found to deviate significantly. Hence, the comparative analysis of real networks with corresponding random networks indicated that they indeed possess some intrinsic properties leading to their diseased and healthy behavior.

The degree distribution plots of all eight networks showed that all, including tumor, adjacent normal, and normal networks, follow the power law of degree distribution, which is an intrinsic feature of true biological networks (Fig. 2A – G). Next, the degree-betweenness and degree-correlation coefficient plots of all eight networks (Fig. 3A – G and Fig. 4A – G) revealed that these networks follow positive and negative correlations, respectively. Several other biological networks have also exhibited the same properties of degree-betweenness and degree-clustering coefficient correlations.

**Fig 2.**
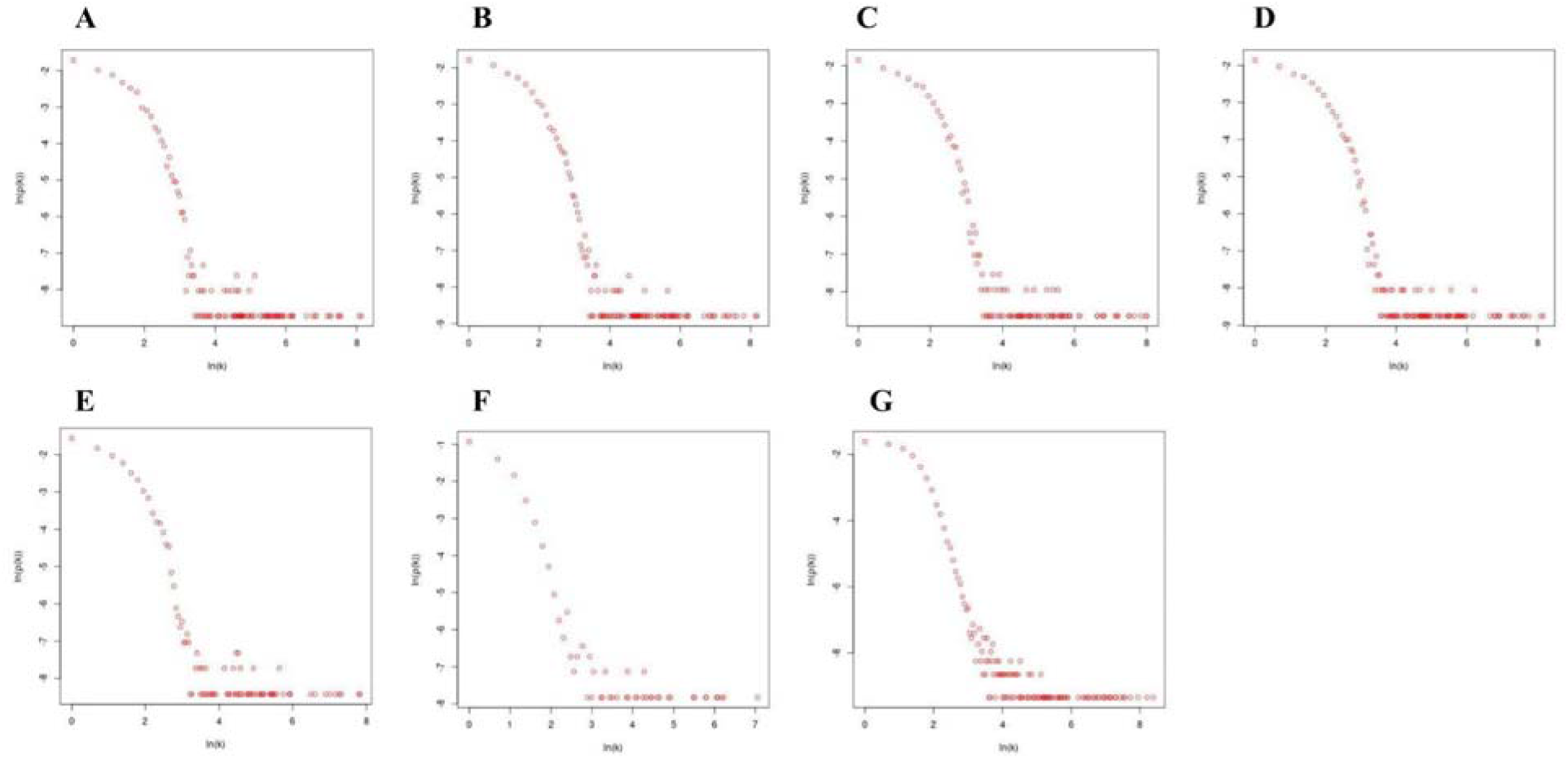
Degree distribution plot of (A) Basal, (B) Her2, (C) LumA, (D) LumB, (E) NormL, (F) ANT, and (G) NBT network

**Fig 3.**
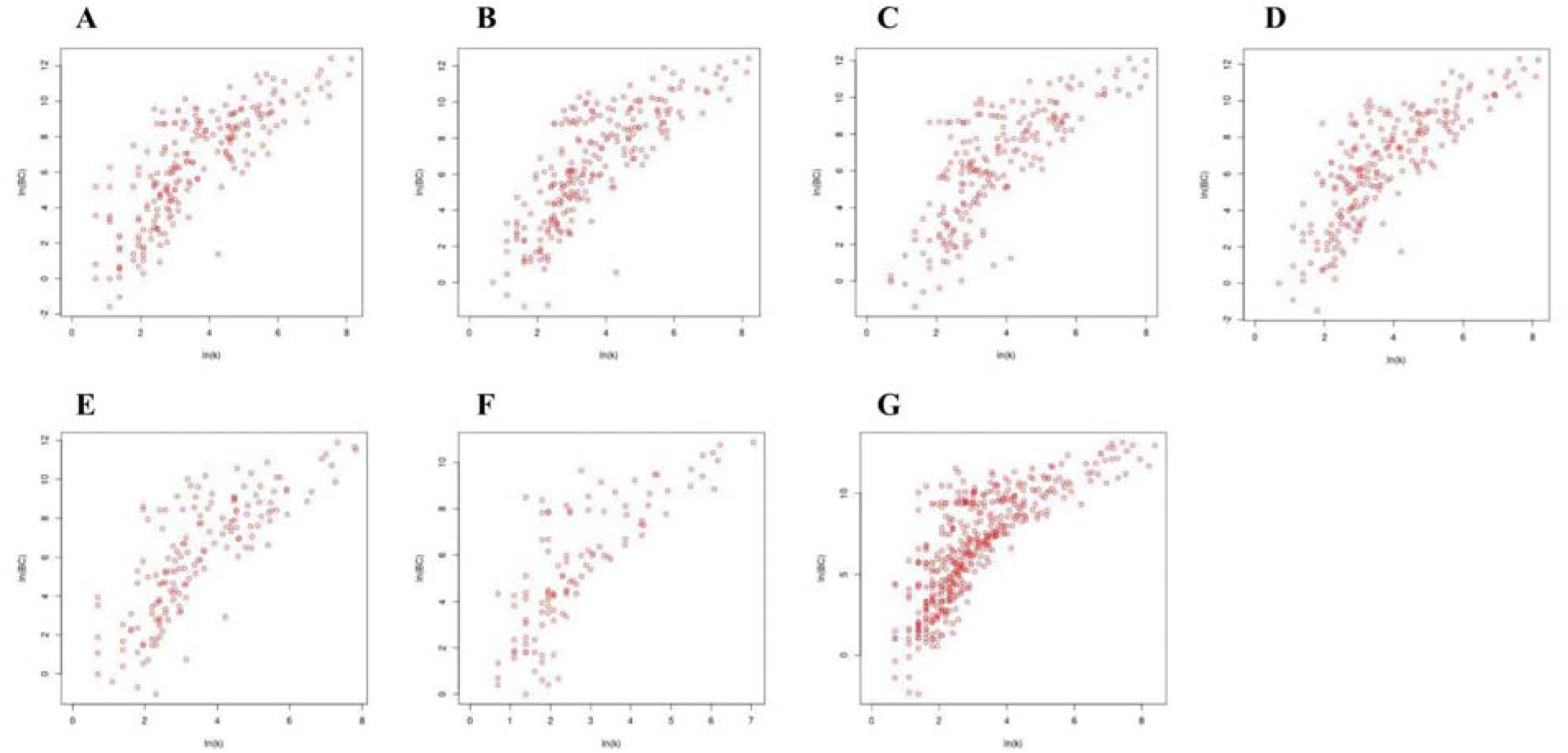
Degree-Betweenness centrality plot of (A) Basal, (B) Her2, (C) LumA, (D) LumB, (E) NormL, (F) ANT, and (G) NBT network

**Fig 4.**
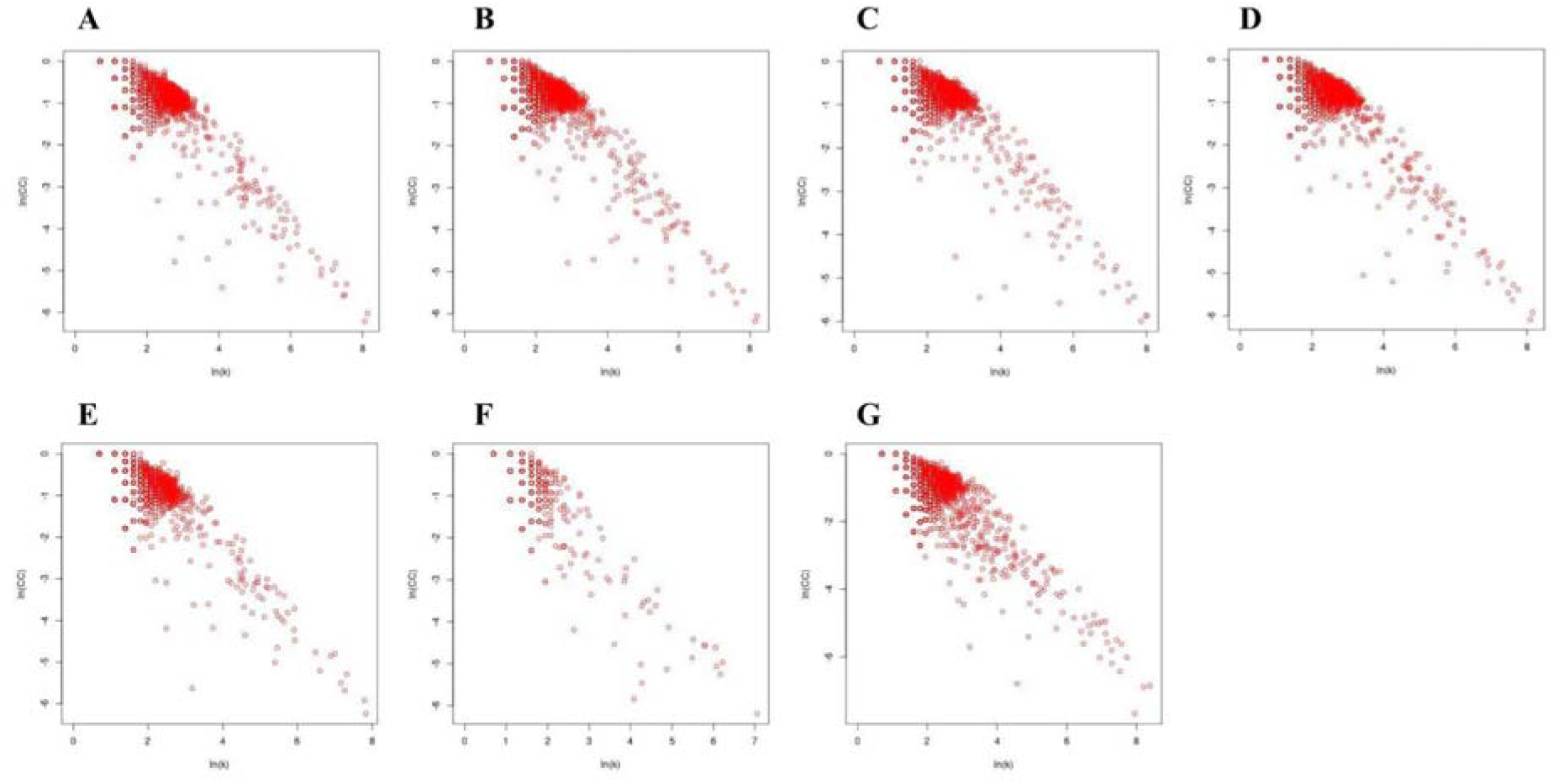
Degree-Clustering coefficient plot of (A) Basal, (B) Her2, (C) LumA, (D) LumB, (E) NormL, (F) ANT, and (G) NBT network

### Similarity of the tumor, adjacent, and normal tissue networks

These networks were also analyzed for their similarity and dissimilarity with each other using the *Jaccard* index based on nodes and edges present therein. Figure 5 displays the similarity plot for all networks based on node and edge *Jaccard* index. Thus, it was observed that the ANT network shared more similarity to the five subtype (tumor) networks as compared to the two normal networks (NBT1 and NBT2).

**Fig 5.**
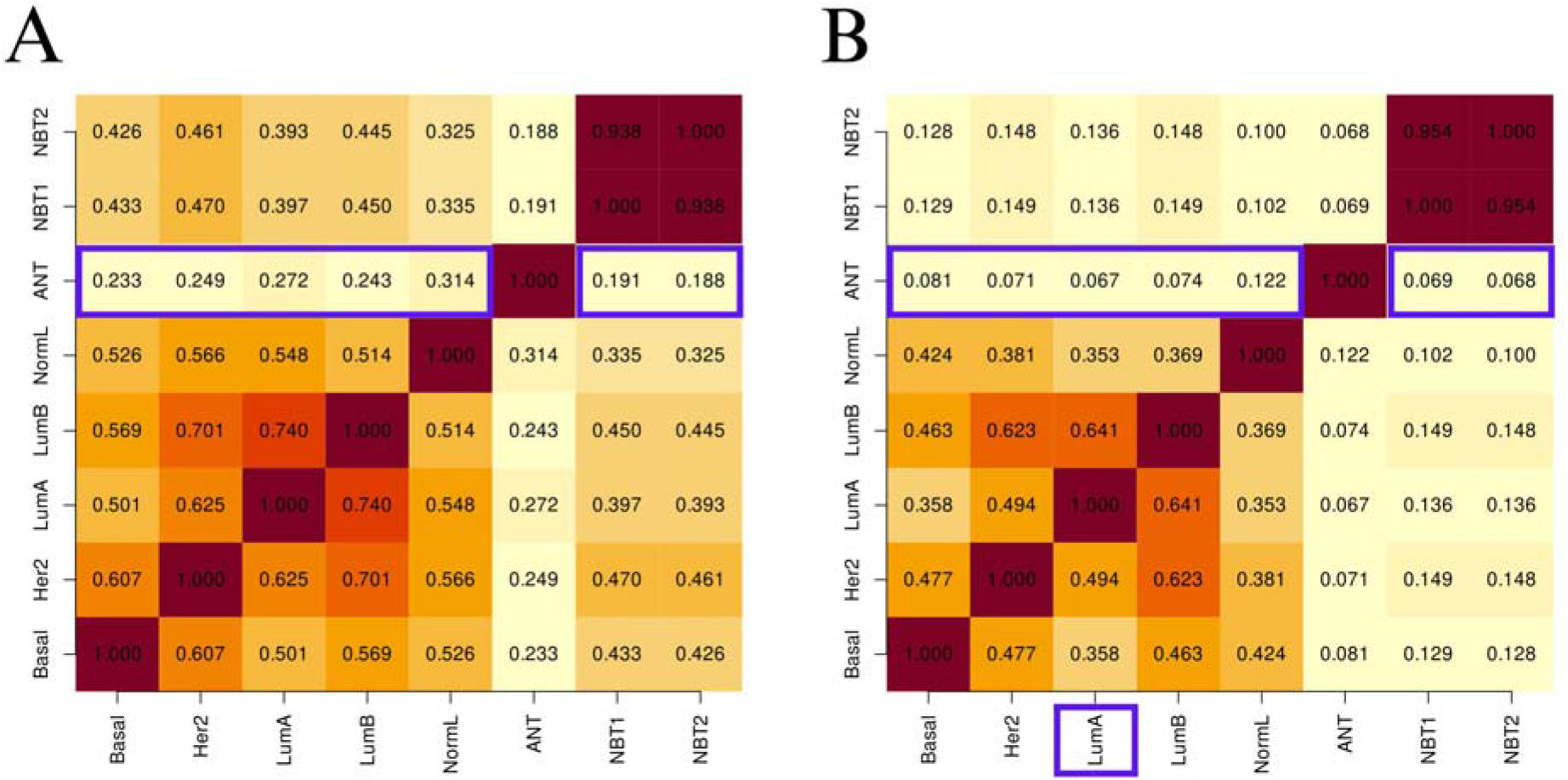
Similarity index (Jaccard) plot for all the tumor, adjacent, and normal tissue networks, (A) Node based *J*- index, and (B) Edge based *J*-index. Red color represents index value close to 1 and white color represents index value close to 0

### Influential genes across tissue types

All subtype and ANT networks were analyzed using EVC+, and key influential genes were identified. Based on the 80:20 principle, mean values of EVC+ of the top twenty percent genes were considered as cutoff values to determine influential genes across different tissue types. The distribution of the total number and tissue-specific number of influential genes are available in Tables 5. There were 24 influential genes that were commonly present in all tissue types, *viz.*, Basal, Her2, LumA, LumB, NormL, and ANT. Among these common influential genes, genes such as *E2F1, FOXA1, JUN, AR, GATA3, BRCA1,* and *ERBB2* were influential among the top key genes based on the value of EVC+ across all subtype networks. Whereas *ESR1* was present only in Basal, LumA, LumB, NormL, and ANT, but not in Her2. Similarly, *TFAP2A* was present in four subtypes, *viz.* Basal, Her2, LumA, and LumB, but not in NormL.

**Table 5.** Distribution of influential genes across different subtypes and ANT.

| Network | Total Number of Influential Genes | Specific Influential Genes | Common Influential Genes |
| --- | --- | --- | --- |
| Basal | 237 | 18 (*ATP6V0A1, BCAR3, CAMTA1, DAPK1, ECE1, ELF1, ELF2, FOXP1, HNRNPL, IGF2BP3, MAD1L1, MEF2D, NFKB2, NUMA1, PHF12, PRMT1, RABGAP1L, SPIB*) | 24 (*E2F1, FOXA1, JUN, AR, KLF4, GATA3, STAT1, STAT2, NR2C2, KLF16, BRCA1, MAZ, MZF1, TCF7L2, CEBPA, SREBF2, GRHL2, TAL1, BCL6, DDIT3, ERBB3, CDH1, CCND1, ERBB2*) |
| Her2 | 254 | 6 (*ARNT, CALM1, MED24, MYO1B, NFIA, SMAD7*) |  |
| LumA | 206 | 5 (*AKAP1, MYO18A, PBX3, SSBP3, TFAP2C*) |  |
| LumB | 229 | 5 (*AHR, COL4A2, PTK2B, REL, TP73*) |  |
| NormL | 145 | 2 (*IRF8, RUNX1*) |  |
| ANT | 78 | 7 (*APC, BMPR2, NFIC, TNS3, USF1, ZBTB7A, ZNF436*) |  |

### Differential correlation analysis and switching gene pairs

After differential correlation analysis of each subtype and adjacent normal tissues with normal tissues at FDR 0.05, we identified the significant number of switching gene pairs present in subtypes and ANT in contrast to normal tissues (Table S2 - S7). We identified a total of 525 switching gene pairs involving 380 genes in the basal subtype (Fig. 6A). In the case of Her2, a total of 428 switching gene pairs involving 465 genes were found (Fig. 6B). Next, the LumA subtype had a total of 176 switching gene pairs comprising 175 genes (Fig. 6C). Further, the LumB contained 186 switching gene pairs involving 206 genes (Fig. 6D). Similarly, the NormL subtype had 813 switching gene pairs involving 677 genes (Fig. 6E). The ANT also exhibited 105 switching genes which involved 120 genes (Fig. 6F). Significant switching gene pairs across subtypes and ANT at correlation coefficients (r1 and r2) 0.3 and FDR 0.05 are given in Table 6.

**Fig 6.**
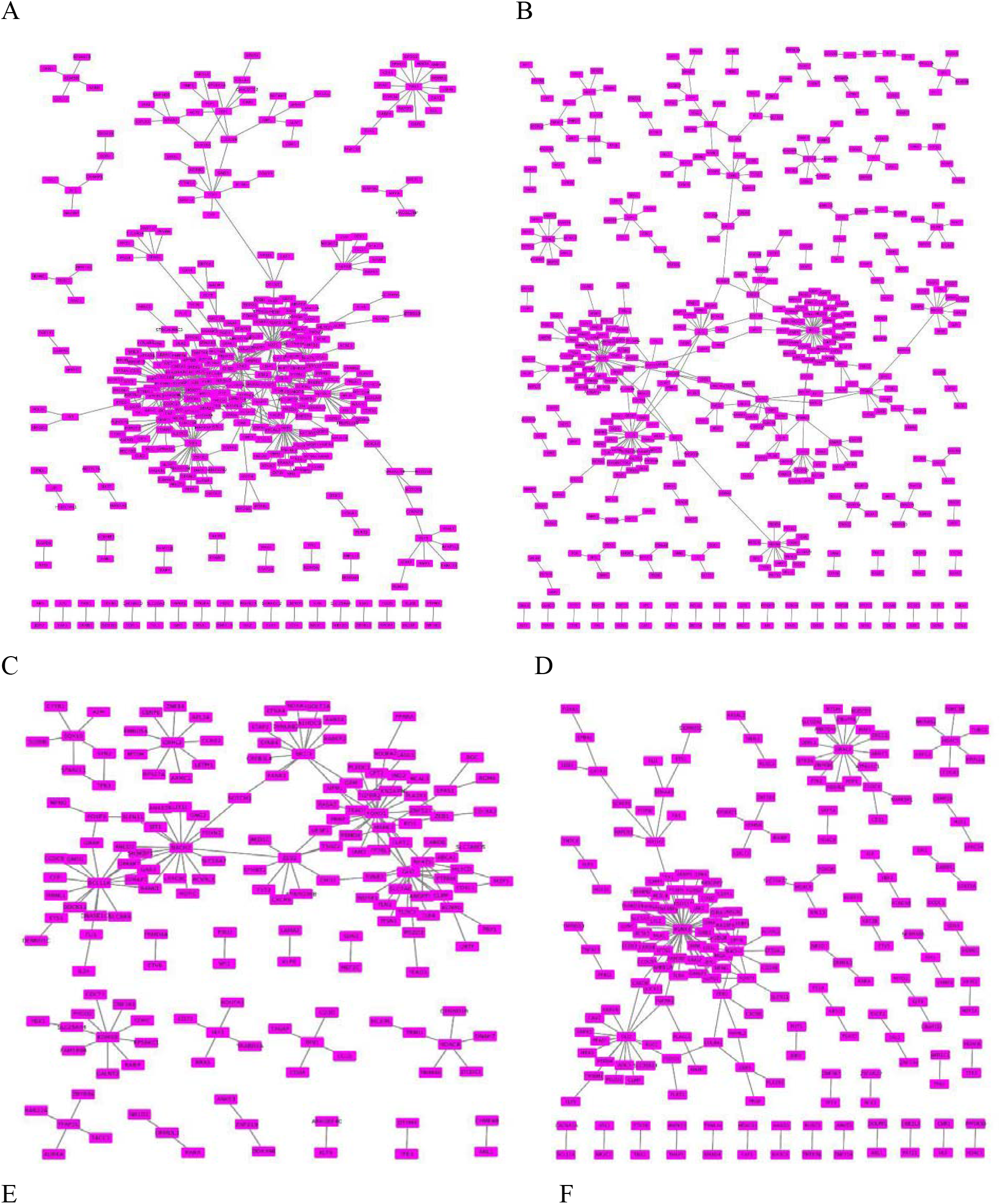
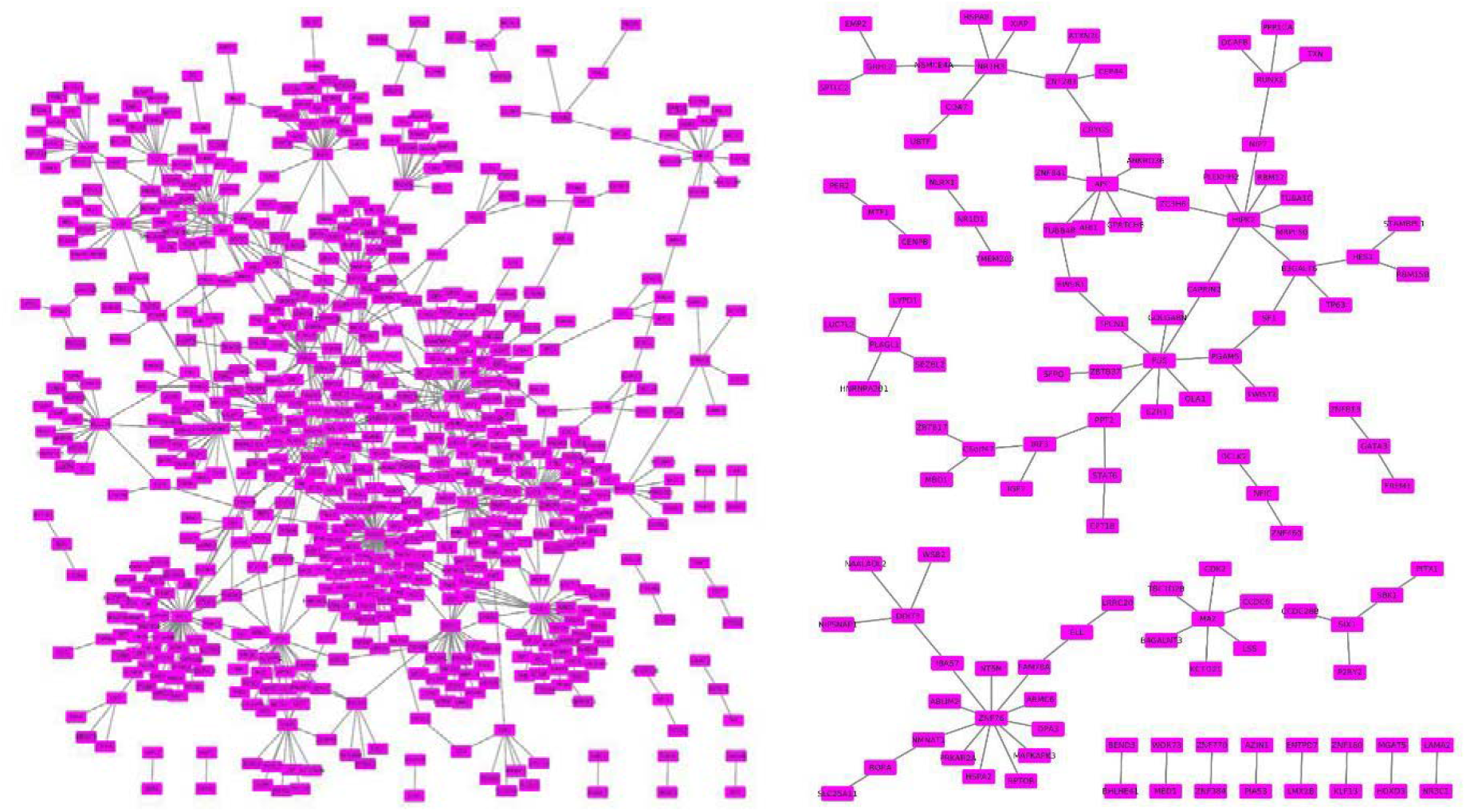
Switching gene regulatory networks of (A) Basal tissues (B) Her2, (C) LumA, (D) LumB, (E) NormL, and (F) ANT

**Table 6.**
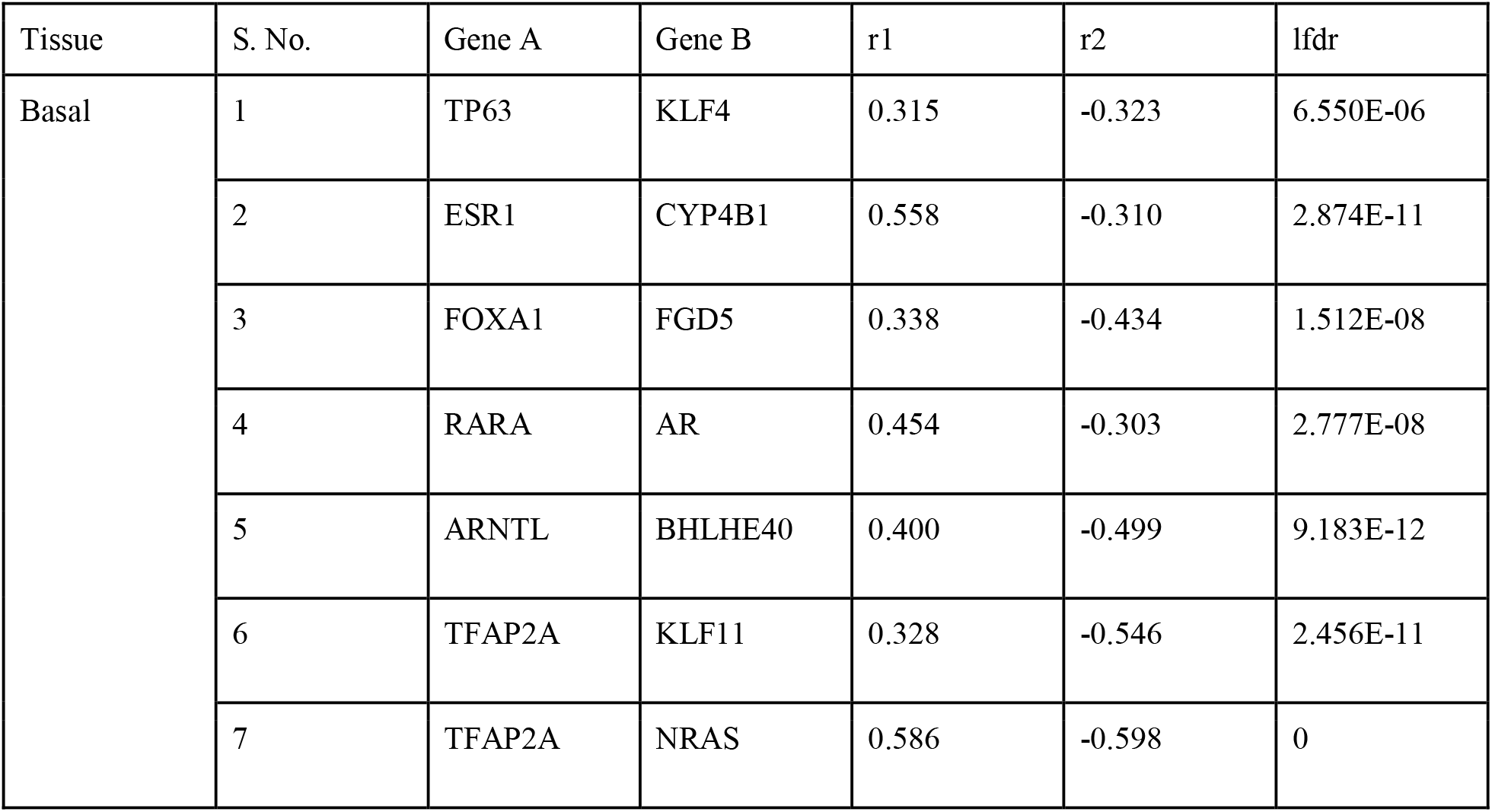
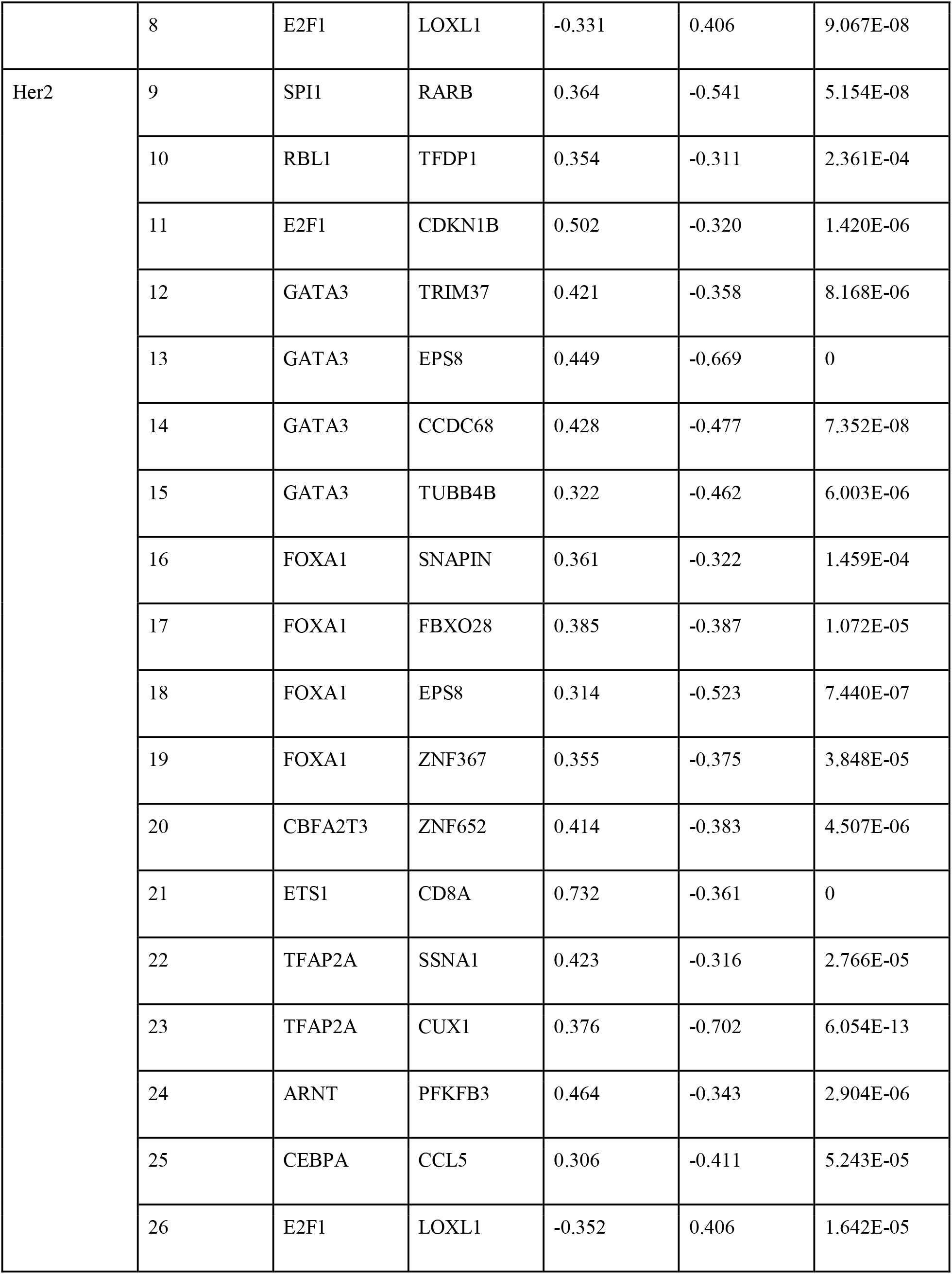
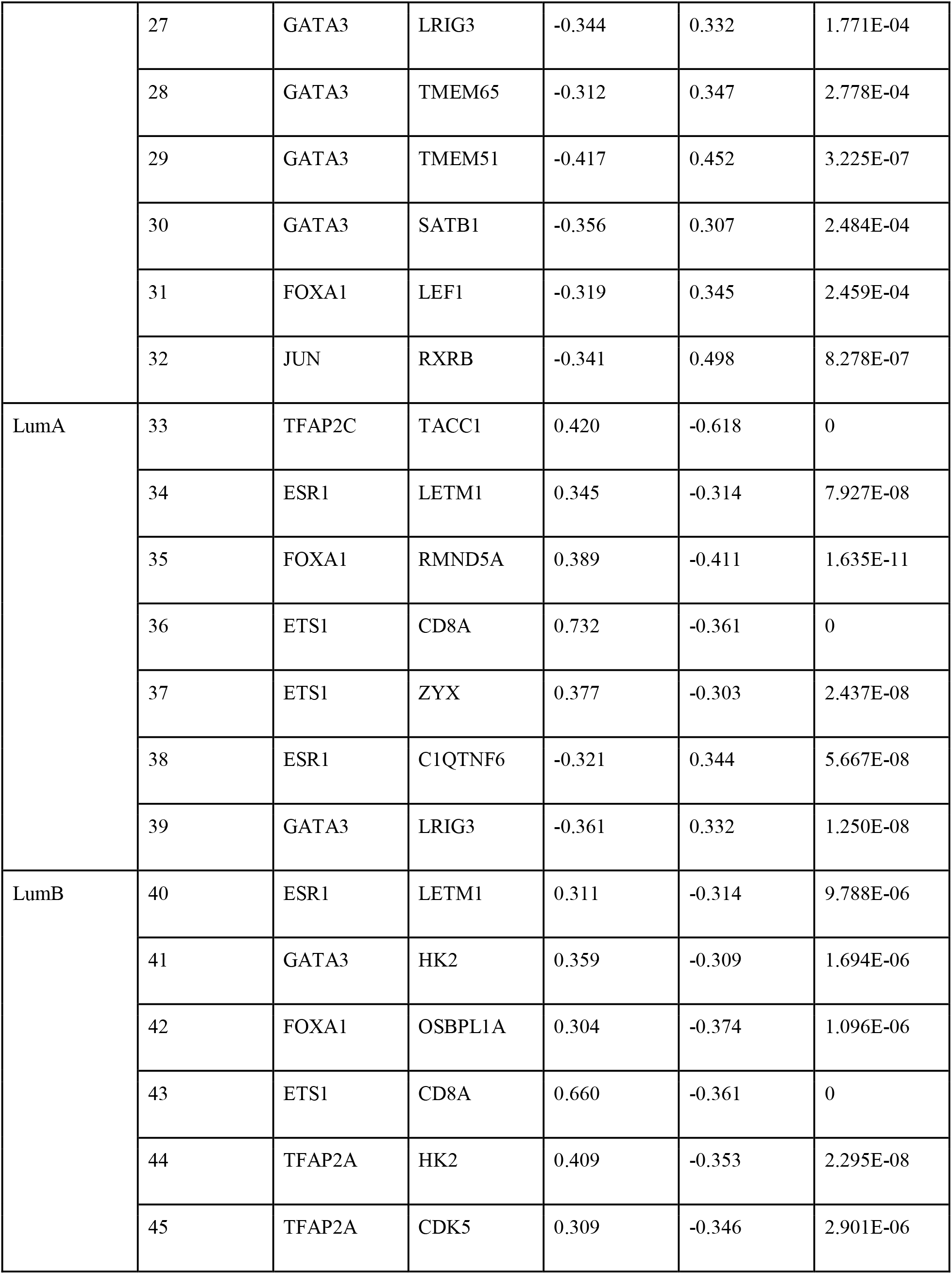
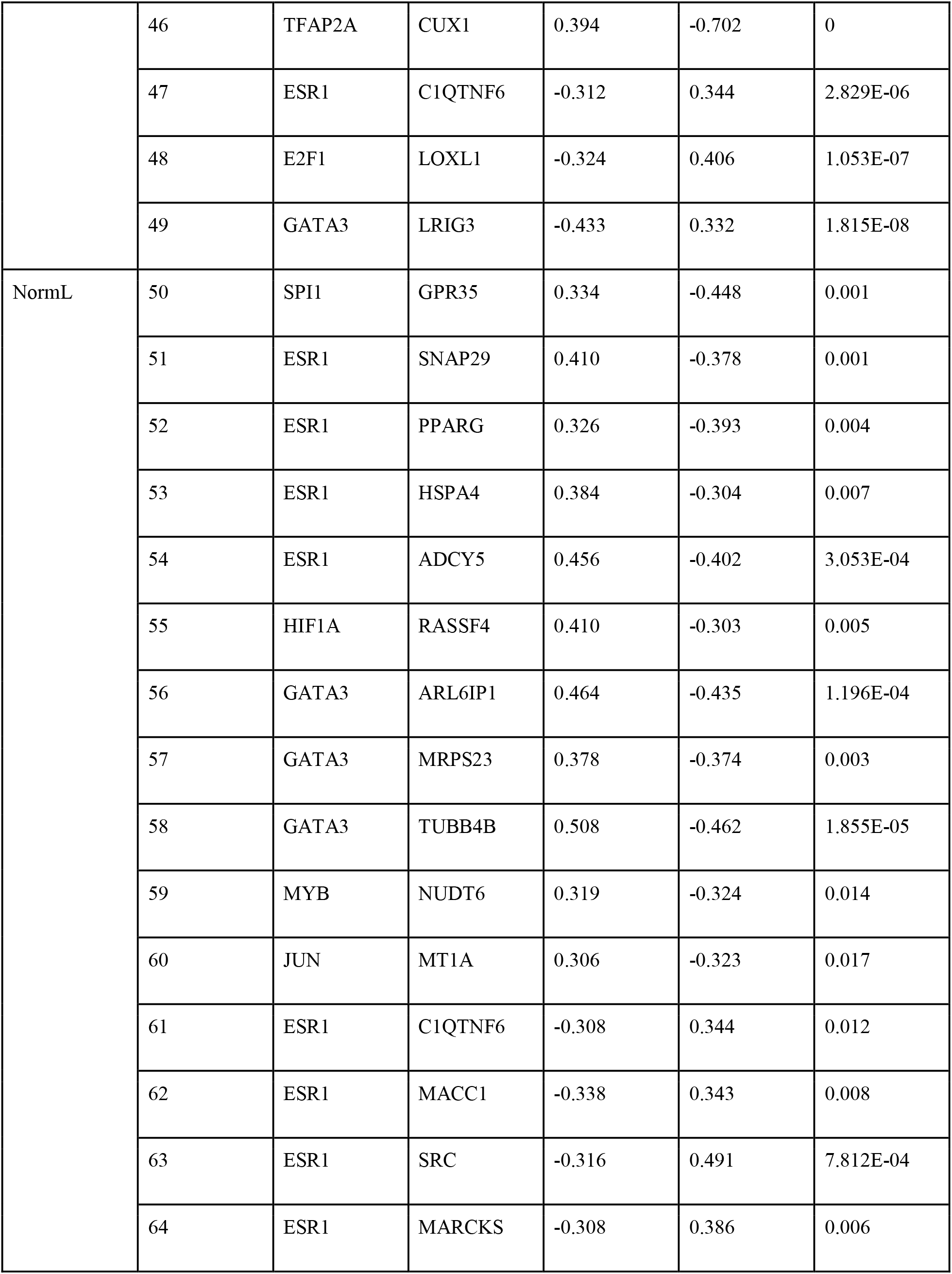
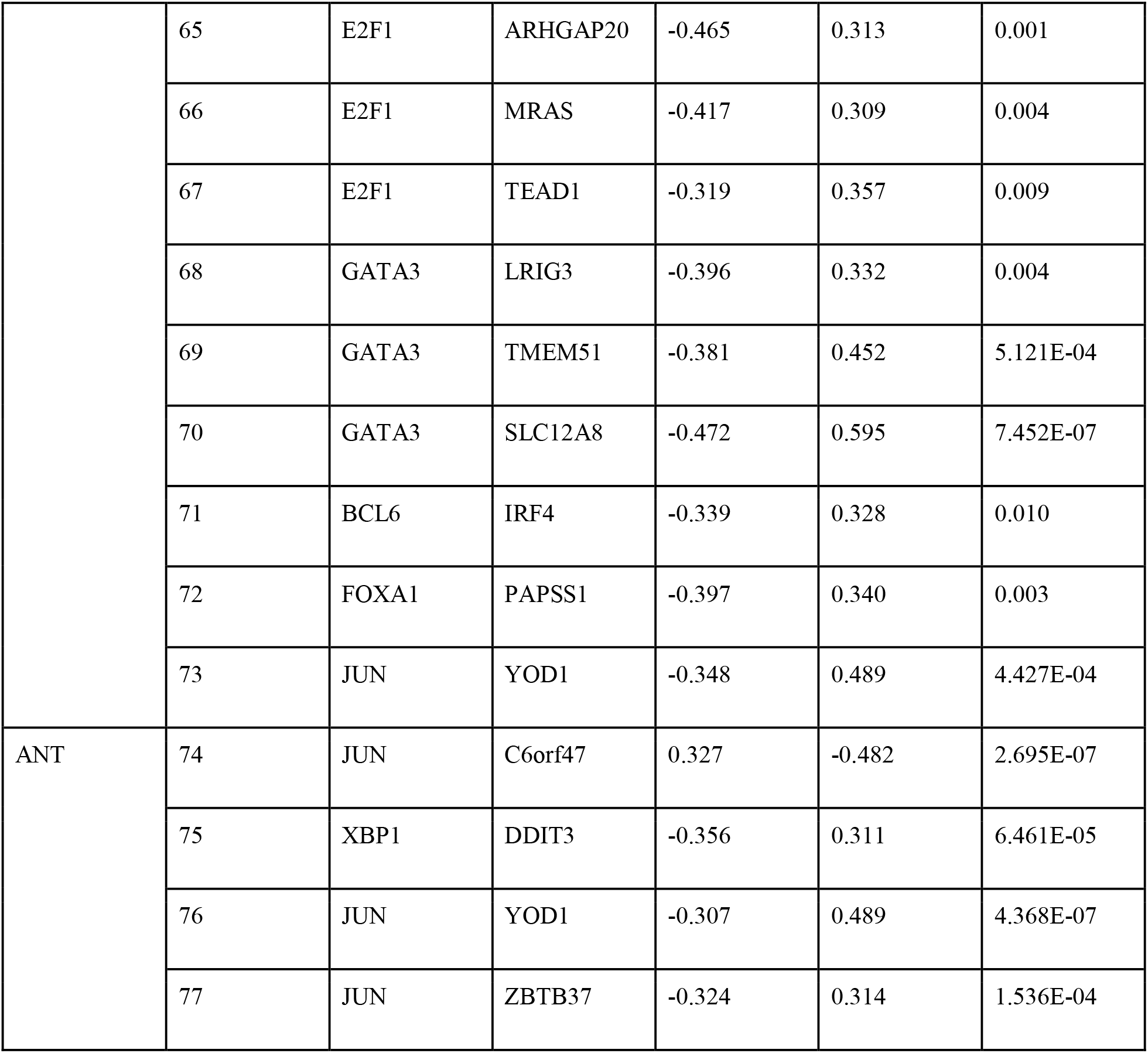
Differentially correlated gene pairs with switching behaviors across six different tissue types at correlation coefficients (r1 and r2) 0.3 and local FDR < 0.05.

### Functional annotations of DEGs

The DEGs identified across all five subtypes were subjected to functional and pathway enrichment analysis to infer the biological processes (BPs) these genes are participating in, the molecular functions (MFs) these genes perform, and the pathways they are involved in. The details of the number of significant GO terms (*viz.* BP and MF) and KEGG pathway terms are available in Table 7. The dot plots in Figure 7 (A - C) represent the top significantly overrepresented GO (BP and MF) and KEGG pathway terms.

**Fig 7.**
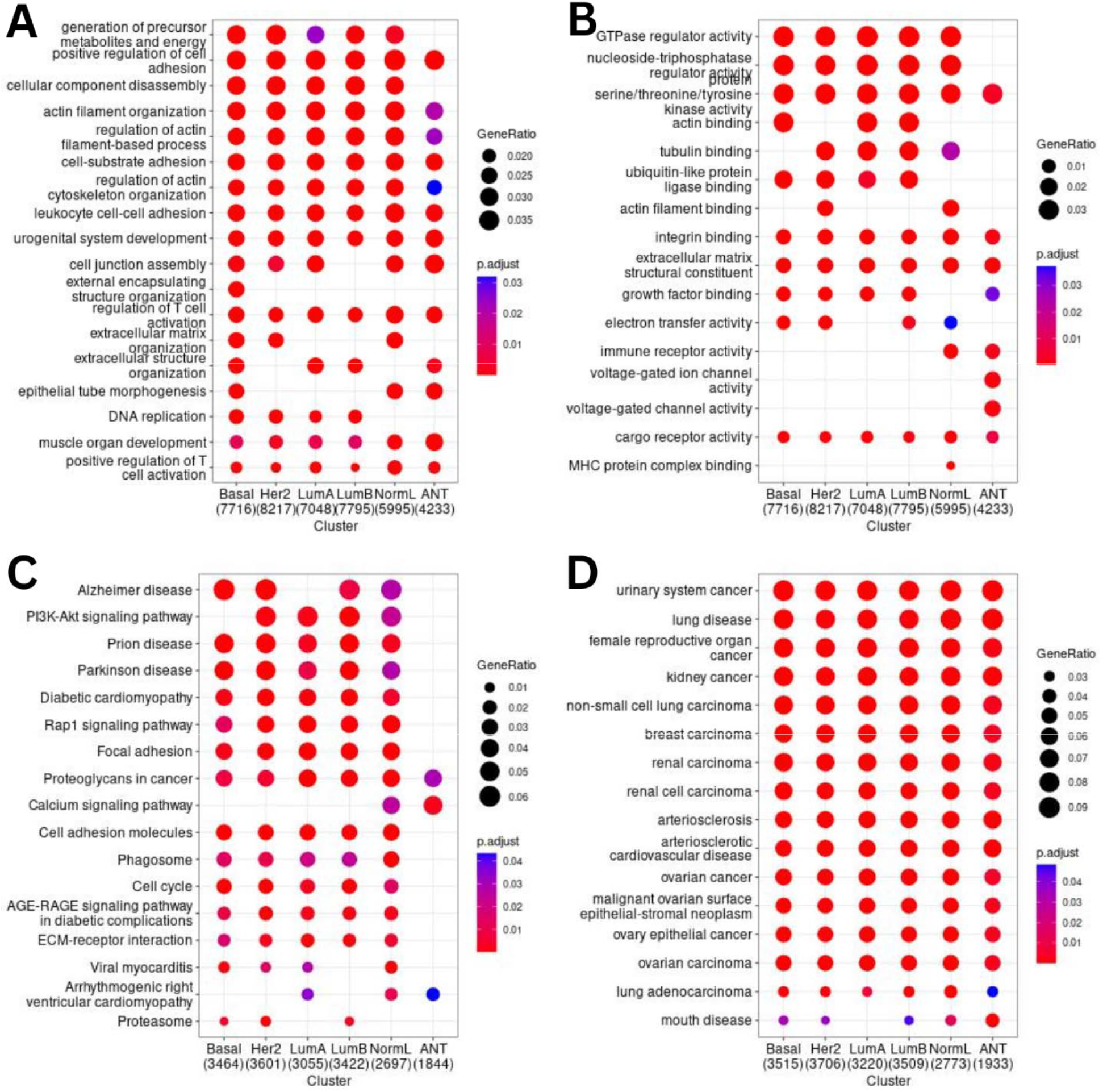
Distribution of top biological process terms (A), molecular functions (B), KEGG pathways (C), and disease ontology (D) based on gene counts associated with DEGs across all five subtypes and ANT

**Table 7.** Distribution of gene ontology, pathway, disease ontology, and disease-gene association terms over-represented among DEGs across breast cancer subtypes and ANT.

| Tissues | GO-BP |  | GO-MF |  | KEGG |  | DO |  | DGN |  |
| --- | --- | --- | --- | --- | --- | --- | --- | --- | --- | --- |
|  | URG | DRG | URG | DRG | URG | DRG | URG | DRG | URG | DRG |
| Basal | 278 | 307 | 54 | 36 | 59 | 31 | 139 | 64 | 1900 | 1086 |
| Her2 | 267 | 313 | 54 | 29 | 69 | 47 | 78 | 89 | 1567 | 1646 |
| LumA | 223 | 317 | 18 | 29 | 48 | 38 | 70 | 108 | 1339 | 1446 |
| LumB | 216 | 316 | 41 | 39 | 44 | 43 | 50 | 95 | 1165 | 1544 |
| NormL | 363 | 192 | 44 | 10 | 83 | 37 | 200 | 29 | 2498 | 623 |
| ANT | 320 | 5 | 23 | 5 | 27 | 2 | 119 | 0 | 2045 | 16 |

Gene ontology - Biological process, GO-BP; Gene ontology - Molecular function, GO-MF; Kyoto encyclopedia of genes and genomes pathway, KEGG; Disease ontology, DO; Disease-gene association network, DGN; Up-regulated gene, URG; and Down-regulated gene, DRG.

Biological processes: There were a total of 1667 and 1450 significant BP terms associated with up-regulated and down-regulated genes, respectively, across Basal, Her2, LumA, LumB, NormL, and ANT (Table 7). Among these enriched terms, BP terms like skeletal system development, external encapsulating structure organization, extracellular matrix organization, extracellular structure organization, positive regulation of cell adhesion, cell-substrate adhesion, leukocyte cell-cell adhesion, urogenital system development, and renal system development were the most significantly enriched terms across all five subtypes and ANT (Fig. 7A, Table S8).

Molecular functions: There were a total of 234 and 148 MF terms associated significantly with up-regulated and down-regulated genes, respectively, across all five subtypes and ANT (Table 7). Of these enriched terms, MF terms like actin binding, glycosaminoglycan binding, extracellular matrix structural constituents, heparin binding, integrin binding, growth factor binding, and transmembrane receptor protein kinase activity were among the significantly enriched terms with a high number of genes as compared to others (Fig. 7B, Table S9).

KEGG pathways: There were a total of 330 and 198 KEGG pathway terms significantly enriched among all up-regulated and down-regulated genes, respectively, across all five subtypes (Basal, Her2, LumA, LumB, and NormL) and ANT. Among all enriched pathway terms, NormL had the highest number of terms. While ANT had the least number of enriched pathway terms (Tables 7 and S10). Among all significantly enriched KEGG pathways terms, four pathways viz. Phagosome, Cell adhesion molecules, Viral myocarditis, and Antigen processing and presentation were associated with up-regulated genes of all subtypes along with ANT. Similarly, two pathways viz. Glycerophospholipid metabolism and Calcium signaling pathway were associated with down-regulated genes of all subtypes along with ANT (Fig. 7C, Table S10).

### Disease ontology and disease-gene associations

The DEGs across all five different subtypes and ANT of breast cancer were subjected to the enrichment of Disease Ontology (DO) terms using the clusterProfiler. Further, these DEGs were also looked for their associations with various diseases using the disease-gene network (DGN) association functionality of clusterProfiler based on the DisGENET database. There were 656 and 385 DO terms significantly enriched with all up-regulated and down-regulated genes, respectively, across all subtypes and ANT. Similarly, there were 10514 and 6361 DGN terms significantly associated with all up-regulated and down-regulated genes, respectively, across different subtypes and ANT (Table 7).

Disease ontology enrichment of DEGs showed significant associations of most of them with several cancers like urinary system cancer, female reproductive organ cancer, kidney cancer, lung cancer, breast cancer, and ovarian cancer (Fig. 7D and Table S11). Similarly, the Disease-Gene association analysis also showed significant associations of these DEGs with cancers like Non-infiltrating intraductal carcinoma, Invasive carcinoma of breast, Metastatic malignant neoplasm to brain, Basal-like breast carcinoma, and Anaplastic carcinoma. Some other diseases, e.g., Infection, Hepatitis C, and Adult meningioma, were also found to be significantly associated with DEGs of different subtypes and ANT (Table S12).

## Discussion

The present study explores comprehensive gene regulatory networks (GRN) of tumor and normal tissues of the female breast to uncover tumorigenic features of adjacent normal tissues and identify key novel genes responsible for the pathogenesis of breast cancer. While the tumor tissues involve five different subtypes of breast cancer, the normal tissues involved morphologically normal adjacent tissues (ANT) and healthy normal tissues (NBT). Each subtype and ANT exhibited different sets of differentially expressed genes (DEGs) and indicated gene expression profile differences among them. The *TP53* was among DEGs across all subtypes but not in the case of ANT. The DEG *MYC* was present in Her2, LumA, and LumB but not in Basal, NormL, and ANT. Some well-known genes of breast cancer, *viz.*, *BRCA1, ESR1*, and *ERBB2,* were too differentially expressed in ANT in addition to all subtypes.

The topological analyses (diameter, betweenness centrality, clustering coefficient, heterogeneity, and average path length) of GRNs of five subtypes, one ANT and one NBT, clearly highlighted the key differences between tumor and normal tissues. The less number of nodes having clustering coefficients’ values zero and one in tumor and adjacent normal tissue compared to normal tissue network indicated that the tumor and adjacent normal networks have similar patterns of wiring among TF and target genes with a decrease in the number of complete subgraphs and terminal nodes. Similar findings have been reported in cervical cancer (Jalan et al. 2015) and breast cancer (Rai et al. 2014), involving protein-protein interaction network-based studies of tumor and normal conditions. Nodes with CC equal to one form complete sub-graphs, and these sub-graphs are the building blocks of a network. Thus, from the above-mentioned result, it can be inferred that the tumor networks are disrupted compared to the normal ones. Similarly, the network of adjacent tissues also has fewer building blocks and hence, is disrupted. Recent studies have indicated that adjacent normal tissues of tumors are not entirely normal. Though it is morphologically normal tissues, these tissues have some tumor-similar features in terms of expression profiles and wiring among TF-target genes. The current findings, such as DEGs of ANT, and topological properties of the ANT network, indicate the same.

Escape velocity centrality-based influential genes prediction led to the identification of 24 key genes, *viz., E2F1, FOXA1, JUN, AR, KLF4, GATA3, STAT1, STAT2, NR2C2, KLF16, BRCA1, MAZ, MZF1, TCF7L2, CEBPA, SREBF2, GRHL2, TAL1, BCL6, DDIT3, ERBB3, CDH1, CCND1,* and *ERBB2*, which were present in all subtypes as well as ANT. These common influential genes are well-known for their role in the development and progression of various cancers, including breast carcinoma. The *E2F1*, a member of the E2F family, is a transcription factor involved in the G1/S transition; it has been reported to control the expression of numerous pro-metastatic genes and act as a master regulator of metastasis in breast cancer. For example, it drives breast cancer metastasis by regulating the gene *FGF13* and altering cell migration (Hollern et al. 2019). Next, the expression of *FOXA1* correlates with LumA and is a significant cancer-specific survival predictor in ER-positive patients (Badve et al. 2007). However, a study by Zhang et al. has confirmed that the overexpression of *JUN* leads to a more invasive phenotype in breast cancer cells and hence, is involved in breast cancer metastasis (Zhang et al. 2007). Further, it has been demonstrated that androgen-induced anti-proliferative effects on *AR*-positive breast cancer cells are *AR*-mediated (Szelei et al. 1997). The *GATA3* (GATA binding protein 3) gene is a transcriptional activator, and a study by Usary et al. suggests that loss of *GATA3* may lead to tumorigenesis in ER+ breast cancers (Usary et al. 2004). However, the *BRCA1* gene, a tumor suppressor with binding activity associated with DNA damage response, has been found among mutated candidate genes in breast cancer (Sjoblom et al. 2006). Similarly, the *ERBB2* (Human epidermal growth factor receptor 2) gene has involvement in breast cancer progression and is evident from the studies reporting that the overexpression of *ERBB2* leads to increased metastasis of breast cancer (Tan et al. 1997; Moody et al. 2002; Holbro et al. 2003).

The differential correlation analysis and uncovering of switching gene pairs across six different tissue types (*viz.* Basal, Her2, LumA, LumB, NormL, and ANT) compared to NBT revealed the essential differences in regulatory patterns or wiring among TFs and target genes acting as switching entities with opposite expressional behavior between tumor/adjacent normal and normal tissues. Many well-known breast cancer genes were significantly differentially correlated with switching behavior across subtypes and ANT. The TF-gene pairs *ESR1, GATA3*, *NFKB2*, and *JAK2* were found to be positively differentially correlated in the Basal subtype while negatively differentially correlated in the case of normal tissues.

The functional enrichment and pathway analysis indicated that the DEGs across subtypes and ANT were significantly associated with cancer-related terms such as cell-cell adhesion, DNA repair, calcium metabolism, and cell cycle. The disease ontology (DO) terms significantly enriched with up-regulated genes across subtypes and ANT included various types of cancers, including breast, renal, ovarian, and lung. The comparative analysis of these terms across subtypes and ANT indicated their similarity and differences. For example, the KEGG pathway term “Proteoglycans in cancer” was present across all tissue types, including different subtypes and ANT, while the term “Calcium signaling pathway” was present only in NormL and ANT. Further, as reviewed by Harjunpää and team, the malignant cells may utilize the pathway of cell adhesion molecules to advance tumor growth and metastasis (Harjunpää et al. 2019). In the present study, the pathway of cell adhesion molecules was found significantly enriched with up-regulated genes across all subtypes and ANT. Among all enriched disease-gene (DGN) terms, few DGN terms like Myeloproliferative disease were found significantly enriched with both up-regulated and down-regulated genes across all subtypes along with ANT. Further, many DGN terms were significantly associated with both up-regulated and down-regulated genes across all subypes except NormL and also were not enriched in case of ANT; for example, terms like Noninfiltrating Intraductal Carcinoma and Invasive carcinoma of breast were associated with all subtypes except the NormL subtype and ANT. Here, it is interesting to note that few cases of myeloproliferative disease-breast cancer associations have been reported (Cornfield et al. 1977).

## Conclusions

The present framework of disease exploration using network biology provides a time and cost-effective approach for understanding the disease complexity. The gene regulatory networks of different subtypes and ANT coupled with differential correlation analysis detected structural and expressional patterns of genes responsible for the disease’s onset and progression. Further, the network analysis using EVC+ helped to identify 24 common and functionally significant genes, including well-known cancer genes such as *E2F1, FOXA1, AR, JUN, BRCA1, GATA3,* and *ERBB2* across subtypes and ANT. Similarly, the EVC+ also helped us to identify tissue-specific genes (Basal: 18, Her2: 6, LuminalA: 5, LuminalB: 5, Normal-Like: 2, and ANT: 7). Moreover, the differential correlation and functional and disease annotation also highlighted the cancer-associated role of these genes. Thus, these common (24) and tissue-specific key genes, especially ANT-specific seven genes, *viz., APC, BMPR2, NFIC, TNS3, USF1, ZBTB7A,* and *ZNF436,* could be utilized for the development of early-stage-specific efficient diagnostic and therapeutic candidates. Further, this analysis framework could also be helpful for other cancer types and diseases, e.g., lung cancer, cervical cancer, head and neck cancer, neuronal disorders, and cardiovascular diseases, towards an understanding of the disease by the disease state’s structural and functional aspects.

## Data availability

The datasets analyzed during the current study are available in public repositories such as The Cancer Genome Atlas (TCGA) and Genotype Tissue Expression (GTEx). Further, The original contributions and/or datasets presented in the study are included in the article/supplementary material.

## Acknowledgments

The authors would like to thank the Center for Modeling, Simulation & Design (CMSD), University of Hyderabad, for providing computational facilities. Further, VV would like to thank the Indian Council of Medical Research (ICMR), New Delhi (ISRM/12(72)/2020, ID: 2020- 2951), Institution of Eminence – University of Hyderabad (No. UoH/IoE/RC3-21-052), and Department of Biotechnology (DBT) – Government of India, New Delhi (No. BUILDER-DBT-BT/INF/22/SP41176/2020) for their financial support. SK also acknowledges the University of Hyderabad for Non-NET Fellowship and ICMR for Senior Research Fellowship (Grant No.: 3/2/2/113/2019/NCD-III, ID: 2019-6723).

## Declarations

### Ethics approval and consent to participate

Not applicable.

### Consent to publish

Not applicable.

### Author contributions

SK and VV conceived and designed the study. SK performed the experiments and analyzed the data. SK drafted, wrote, and edited the manuscript. VV reviewed and edited the manuscript. VV supervised the study. All authors read and approved the final manuscript.

## Supplementary Tables

Table S1. Topological properties of random networks.

Table S2. Switching gene pairs of Basal subtypes with adjP-value < 0.05 and |Correlation coefficients| > 0.4.

Table S3. Switching gene pairs of Her2 subtypes with adjP-value < 0.05 and |Correlation coefficients| > 0.4.

Table S4. Switching gene pairs of LumA subtypes with adjP-value < 0.05 and |Correlation coefficients| > 0.4.

Table S5. Switching gene pairs of LumB subtypes with adjP-value < 0.05 and |Correlation coefficients| > 0.4.

Table S6. Switching gene pairs of NormL subtypes with adjP-value < 0.05 and |Correlation coefficients| > 0.4.

Table S7. Switching gene pairs of ANT with adjP-value < 0.05 and |Correlation coefficients| > 0.4.

Table S8. GO-BP terms significantly associated with DEGs at adjP-value < 0.05. Table S9. GO-MF terms significantly associated with DEGs at adjP-value < 0.05.

Table S10. KEGG pathway terms significantly associated with DEGs at adjP-value < 0.05.

Table S11. Disease ontology terms significantly associated with DEGs at adjP-value < 0.05.

Table S12. Disease-gene network terms significantly associated with DEGs at adjP-value < 0.05.

