## Supplementary Table S1 for "Architecture and topologies of gene regulatory networks associated with breast cancer, adjacent normal, and normal tissues"

Table S1. Topological properties of seven random networks corresponding to real networks.

| **Network** | **<k>** | **<BC>** | **<EC>** | **<CC>** | **CC=1** | **CC=0** | **APL** | **D** | **HC** |
| --- | --- | --- | --- | --- | --- | --- | --- | --- | --- |
| Basal | 10.783 | 134.094 | 0.020 | 0.733 | 964 | 491 | 2.741 | 6 | 7.855 |
| Her2 | 11.055 | 118.045 | 0.021 | 0.675 | 610 | 815 | 2.677 | 6 | 8.098 |
| LumA | 11.683 | 90.786 | 0.024 | 0.829 | 1090 | 35 | 2.550 | 6 | 7.722 |
| LumB | 11.943 | 100.949 | 0.022 | 0.795 | 1107 | 27 | 2.538 | 6 | 7.742 |
| NormL | 9.022 | 68.464 | 0.023 | 0.672 | 363 | 519 | 2.459 | 6 | 7.831 |
| ANT | 4.645 | 21.515 | 0.020 | 0.858 | 999 | 28 | 2.216 | 4 | 6.936 |
| NBT | 8.018 | 190.695 | 0.013 | 0.585 | 1450 | 1589 | 2.991 | 8 | 10.032 |

Average degree, <k>; Average betweenness centrality, <BC>; Average eigenvector centrality, <EC>; Average clustering coefficient, <CC>; Number of nodes with CC = 1, N (CC=1); Number of nodes with CC = 0, N (CC=0); Average path length, APL; Diameter, D; Heterogeneity centrality, HC.
